# Alleviating chronic ER stress by p38-Ire1-Xbp1 pathway and insulin-associated autophagy in *C. elegans* neurons

**DOI:** 10.1101/2020.03.02.972794

**Authors:** Liying Guan, Zhigao Zhan, Yongzhi Yang, Yue Miao, Xun Huang, Mei Ding

## Abstract

ER stress occurs in many physiological and pathological conditions. However, how chronic ER stress is alleviated in specific cells in an intact organism is an outstanding question. Here, overexpressing the gap junction protein UNC-9 (Uncoordinated) in *C. elegans* neurons triggers the Ire1-Xbp1-mediated stress response in an age-dependent and cell-autonomous manner. The p38 MAPK PMK-3 acts upon the phosphorylation of IRE-1 to regulate chronic stress. Overexpressing gap junction protein also activates autophagy and the insulin pathway functions through autophagy, but not the transcription of genes encoding ER chaperones, to counteract the p38-Ire1-Xbp1-mediated stress response. Together, these results reveal an intricate cellular regulatory network in response to chronic stress in a subset of cells in multicellular organism.

**Author Summary:** The accumulation of unfolded proteins triggers the ER stress response (UPR), which allows cells to fight against fluctuations in protein expression under both physiological and pathological conditions. Severe acute ER stress responses can be induced by drug treatment. However, such intense ER stress rarely occurs ubiquitously in every cell type *in vivo*. Here, we designed a genetic system in the nematode *C. elegans*, which allows us to induce ER stress in specific cells, without drug treatment or any other external stimuli, and then to monitor the stress response. Using this system, genetic screens identified that the p38 MAPK directly acts through the IRE-1-XBP-1 branch of the UPR to promote the stress response. Meanwhile, the insulin receptor functions through autophagy activation to counteract the p38-IRE-1-XBP-1 pathway. Together, these results reveal an intricate cellular regulatory network in response to chronic stress in multicellular organism.

## Introduction

Proteins destined for secretion or insertion into membranes enter the endoplasmic reticulum (ER) in an unfolded form and generally leave only after they have reached their native states. During normal function, a secretory cell may experience dramatic variations in the flux of new proteins through its ER in response to changes in demand. Disruption of the balance between secretory protein synthesis and the folding capacity of the ER activates a signaling network called the Unfolded Protein Response (UPR) in an attempt to maintain homeostasis [1]. The UPR reduces protein translation, increases expression of ER chaperones and enzymes to facilitate protein folding, and clears misfolded proteins for degradation [1]. Extensive or prolonged UPR activity signals that the accumulation of misfolded proteins has overwhelmed the compensatory mechanisms of the UPR, and an apoptotic response may be elicited [2]. In contrast, certain types of cancer use the protective role of the UPR to sustain their rapid growth [3, 4]. Therefore, the UPR can serve either as an apoptotic executor or as a cytoprotector, depending on the cellular context.

In multicellular organisms, the UPR is comprised of three branches, mediated by the transmembrane ER luminal sensors IRE1 (inositol requiring enzyme 1), PERK (double-stranded RNA-activated protein kinase [PKR]-like ER kinase), and ATF6 (activating transcription factor 6) [1]. Of the three, IRE1 is the only ER-stress sensor protein that is conserved from yeast to human [5]. In response to ER stress, IRE1 oligomerizes to activate an endoribonuclease domain that splices the mRNA encoding the transcription factor HACT1 (homologous to ATF/CREB) in yeast [6, 7] or XBP1 (X-box binding protein 1) in metazoans [8, 9]. The spliced *xbp1* mRNA produces a short, activated form of XBP1, called XBP1s, which activates expression of genes encoding chaperones and other ER-associated degradation proteins that expand the folding capacity of the ER and increase the breakdown of misfolded proteins [10]. In various experimental systems, the induction of ER stress is usually achieved by treatment with drugs, including the glycosylation inhibitor tunicamycin and the ER calcium ATPase inhibitor thapsigargin. It is worth noting, however, that the rapid induction of ER stress by toxins results in a massive and synchronous perturbation of ER function. In contrast, in a variety of human diseases, including Pelizaeus-Merzbacher disease, Huntington’s disease, and osteogenesis imperfecta, the mutant or wild-type forms of secretory proteins gradually accumulate in the ER and cause pathological conditions in a subset of tissues or cell types [11, 12]. *In vivo* models in which misfolded or/and unfolded proteins in the overloaded ER can be directly monitored in limited cell types will certainly help us to explore the molecular mechanisms underlying chronic ER stress responses [13].

Macroautophagy (hereafter referred to as autophagy) is a cellular degradation process initiated in response to stress. It attempts to restore metabolic homeostasis through lysosomal degradation of cytoplasmic organelles or cytosolic components [14]. Autophagy requires a core set of conserved proteins known as autophagy-related (Atg) proteins and is mediated by an organelle called the autophagosome [15]. Autophagosomal membranes originate from the ER, the Golgi complex, mitochondria, endosomes, and the plasma membrane [16, 17]. During formation and expansion of the autophagosome precursor structure, called the phagophore, cytosolic material and damaged subcellular organelles are captured and enclosed. The complete closed autophagosome then fuses with lysosomes to become an autolysosome, inside which degradation and recycling of its content occurs. Autophagy is activated under ER stress conditions and many of the components that mediate autophagy have been identified as UPR target genes and are important for cells to survive severe ER stress [18, 19].

The nematode *C. elegans* is transparent throughout its life cycle. We reasoned that if we could induce the ER stress response by overloading the cell with a non-secreted plasma membrane protein, we would then be able to detect the occurrence of UPR by directly observing the sub-cellular localization of this protein *in vivo*. Gap junction channels direct cell-to-cell communication and defects in gap junction formation and/or function are the cause of at least 10 human pathological conditions [20]. Moreover, the disease-related gap junction proteins accumulate in the ER and trigger the ER stress response [21–24]. Here, we found that overexpressing the gap junction protein UNC-9 in *C. elegans* neurons activates the IRE-1-XBP-1-mediated UPR. Using this model, we further identified that the p38 MAP kinase PMK-3 functions through IRE-1 phosphorylation to regulate UPR. Loss of function of *daf-2* (encoding the insulin-like receptor) suppresses the misfolded/unfolded UNC-9 defect in *pmk-3*, *ire-1* or *xbp-1* mutants. Intriguingly, although the reduced transcription of ER chaperone Bip/*hsp-4* (immunoglobulin heavy chain-binding protein/heat shock protein 4) is not rescued by *daf-2* mutation, autophagy induction is largely restored. This study reveals the intricate molecular interactions underlying the ER stress response in a limited subset of cells in an intact organism, and provides insights into the pathophysiology of many human degenerative diseases.

## Results

### Excess UNC-9::GFP in DD/VD neurons triggers the ER stress response

In *C. elegans*, the D-type motor neurons, including 6 DDs and 13 VDs, are a set of GABAergic neurons with cell bodies distributed along the ventral nerve cord and neuronal processes projecting both along the ventral cord and to the dorsal cord [25]. *unc-25* encodes the GABA biosynthetic enzyme glutamic acid decarboxylase [26]. We used the P*unc-25* promoter to specifically overexpress the GFP (green fluorescent protein)-fused gap junction protein UNC-9 in DD/VD neurons. The ectopically expressed UNC-9::GFP is distributed in a punctate pattern along the neuronal processes and inside the cell body region of DD/VD neurons (Fig 1A and 1B). In contrast, an UNC-9::GFP knock-in (KI) line created using the CRISPR-Cas9 technique displays a punctate pattern on neuronal processes and the cell surface, but not within the cell body region of DD/VDs (Fig 1C). Double staining with markers labeling various subcellular compartments demonstrated very little overlap between the ectopic UNC-9::GFP puncta and the ER marker CYTB-5.1::mCherry or the Golgi marker mCherry::MANS (Fig 1B). Instead, a large proportion of the UNC-9::GFP signal was distributed on the plasma membrane (co-localized with Myr::mCherry), early endosomes (co-localized with mCherry::RAB-5) and late endosomes or lysosomes (co-localized with mCherry::RAB-7) (Fig 1B).

**Fig 1.**
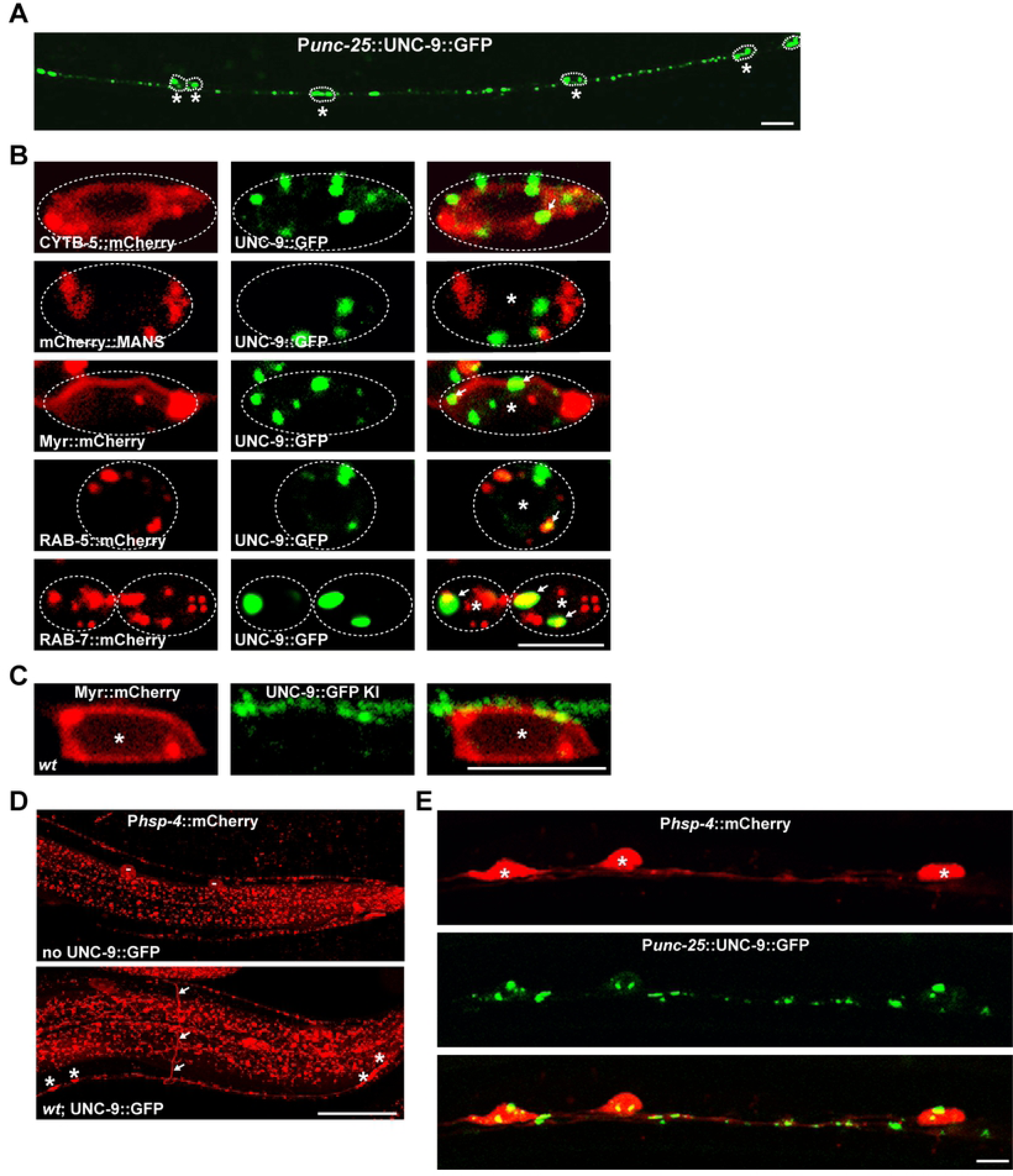
Excess UNC-9::GFP activates the unfolded protein response in DD/VDs. (A) UNC-9::GFP (green) driven by the P*unc-25* promoter is expressed in DD/VD neurons. Individual DD/VD cell bodies are indicated by * and are encircled by dashed lines. Scale bar represents 10 µm. (B) The subcellular localization of UNC-9::GFP (green) in DD/VD cell body regions. The ER, Golgi, plasma membrane, early endosomes and late endosomes/lysosomes are indicated by CYTB-5::mCherry (red), mCherry::MANS (red), Myr::mCherry (red), mCherry::RAB-5 (red) and mCherry::RAB-7 (red) respectively. Individual DD/VD cell bodies are indicated by * and are encircled by dashed lines. (C) UNC-9::GFP, expressed at the endogenous level via knock-in (KI), is distributed on the neuronal processes and cell surface of DD/VD neurons (asterisks). Scale bars in (B) and (C) represent 5 µm. (D) The expression level of P*hsp-4*::mCherry (red) is increased in DD/VDs. White arrows indicate DD/VD commissures. Asterisks indicate DD/VD cell bodies. “-” marks the scavenging cells (coelomocytes) which are inconsistently highlighted by P*hsp-4*::mCherry. Scale bar represents 25 µm. (E) P*hsp-4*::mCherry (red) signal is induced in P*unc-25*::UNC-9::GFP (green) expressing cells. Scale bar represents 5 µm.

To test whether the overexpression of UNC-9 in DD/VDs triggers the ER stress response, we examined the expression of a reporter containing mCherry fused to the promoter of the gene *hsp-4*. *hsp-4* encodes the worm homolog of the ER chaperone protein Bip and its transcription is strongly induced by the stress response [9]. In the absence of ectopic UNC-9::GFP, the P*hsp-4*::mCherry is widely but weakly distributed in multiple tissues (S1A Fig). When UNC-9::GFP was overexpressed in DD/VD neurons, we found that the mCherry signal was not increased in any other neurons or other tissues (S1A and S1B Fig). Instead, P*hsp-4*::mCherry was specifically induced in DD/VD neurons (Fig 1D and 1E). There are more than 60 neurons with their cell bodies distributed along the ventral cord [25]. As shown in Fig 1D, only the commissures (white arrows) and the cell bodies (asterisks) of the 19 DD/VD neurons (18±0.7, N=22) (S1B Fig) could be distinctly visualized by the P*hsp-4*::mCherry signal. Double-labeling analysis further confirmed that the mCherry signal was co-distributed specifically with the P*unc-25* promoter driven UNC-9::GFP expressing cells (S1B Fig and Fig 1E).

### The IRE1-XBP1 UPR branch regulates the subcellular localization of overexpressed UNC-9::GFP in neurons

The stress response triggered by UNC-9::GFP overexpression is likely employed to facilitate the correct folding of excess UNC-9::GFP protein. We suspected that the disruption of components involved in the underlying stress response process would abolish the punctate distribution pattern of UNC-9::GFP in DD/VD neurons. Hence, we conducted a genetic screen for mutants with altered UNC-9::GFP distribution. From this screen, we isolated an *xbp-1* allele, *xd131*. In *xbp-1(xd131)* animals, the UNC-9::GFP signal no longer takes the characteristic punctate form. Instead, it is diffusely distributed in the cell body region of DD/VD neurons. Except for a few small residual dots, there is no GFP signal on the neurites (Fig 2A). Other *xbp-1* alleles display a similar phenotype to *xd131* (Fig 2A). Introducing a wild-type copy of the active but not the inactive *xbp-1* into DD/VD cells significantly rescues the UNC-9::GFP localization defect (Fig 2B), which suggests that loss of function of *xbp-1s* is responsible for the abnormal UNC-9::GFP distribution. *ire-1* mutants show an almost identical phenotype to *xbp-1* mutants (Fig 2A). In contrast, mutation of PERK (*pek-1* mutant) or ATF6 (*atf-6* mutant) did not affect the punctate pattern of UNC-9::GFP in DD/VDs (Fig 2A and S2A Fig). No obvious reduction of UNC-9::GFP expression could be detected in *ire-1*, *xbp-1*, *pek-1* or *atf-6* mutant animals (S2B and S2C Fig).

**Fig 2.**
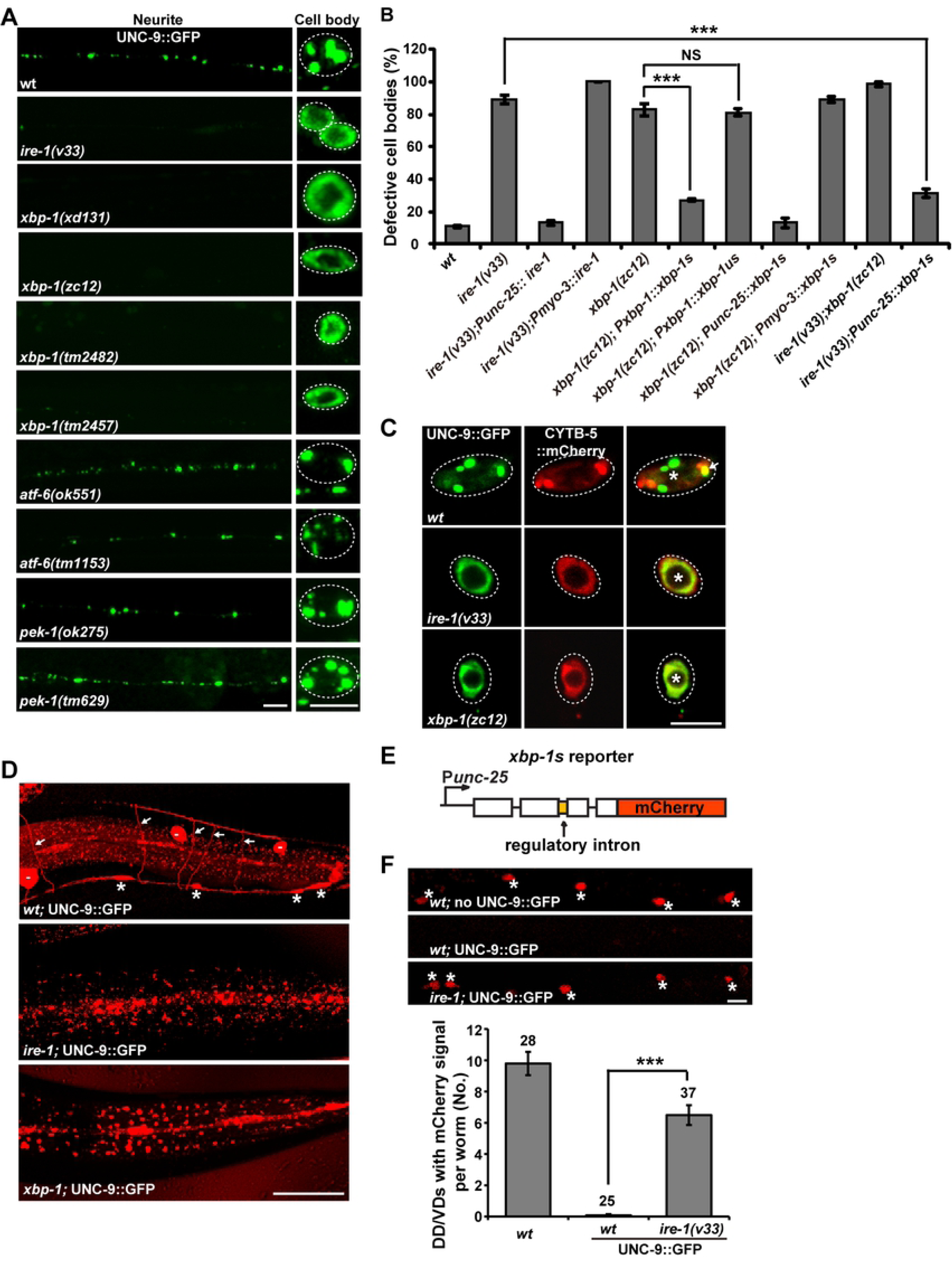
The IRE-1-XBP-1 UPR branch is specifically required for the subcellular localization of excessive UNC-9::GFP. (A) The distribution of UNC-9::GFP (green) driven by the P*unc-25* promoter in various UPR mutants. (B) Quantification of the UNC-9::GFP localization defect in various UPR mutant animals. *N* = 20 for each genotype; \*\*\**P* < 0.001; NS, not significant. One-way ANOVA with Dunnett’s test. (C) Co-localization between UNC-9::GFP (green) and the ER marker CYTB-5::mCherry (red) in *ire-1* and *xbp-1* mutants in the DD/VD cell body region. Scale bars represent 5 µm in (A) and (C). (D) P*hsp-4*::mCherry expression (red) in wild type (*wt*), *ire-1* and *xbp-1* mutants. White arrows indicate DD/VD commissures. Asterisks indicate DD/VD cell bodies. Scale bar represents 25 µm. (E) Schematic drawing of the *xbp-1* reporter, which is specifically expressed in DD/VDs using the P*unc-25* promoter. (F) Assay for the *ire-1*-dependent alternative splicing of *xbp-1* in DD/VDs. Data are expressed as mean ± s.e.m; *N* is indicated above each column; \*\*\**P* < 0.001. One-way ANOVA with Fisher’s Least Significant Difference (LSD) test. Scale bar represents 10 µm.

The *unc-9* gene is also expressed in body wall muscles. *myo-3* encodes the body wall muscle-specific myosin [27, 28]. Using the P*myo-3* promoter, we drove UNC-9::GFP protein expression in body wall muscle cells. In accordance with the observation that the body wall muscle cells are extensively coupled by gap junction connections [29, 30], most of the UNC-9::GFP signal is distributed in a punctate pattern along the muscle edge, particularly at the region where two neighboring muscle cells contact each other (S3A Fig). When the P*myo-3*::UNC-9::GFP marker was introduced into *xbp-1* or *ire-1* mutant animals, however, we found that the UNC-9::GFP puncta largely remained at the cell-cell contact region of muscle cells, similar to wild type (S3A Fig). The different distribution of UNC-9::GFP in DD/VD neurons versus muscle cells in *xbp-1* or *ire-1* mutants implies that *ire-1* and *xbp-1* may not affect the protein localization of UNC-9 in general. Indeed, when we examined the UNC-9::GFP distribution in UNC-9::GFP KI strain created by CRISPR-Cas9, we found that the UNC-9::GFP signal expressed from the KI strain retains its punctate pattern in both *ire-1* and *xbp-1* mutant animals, and is distributed on the neuronal processes and the cell surface of DD/VDs in a pattern that is indistinguishable from wild type (S3B Fig).

The P*unc-53* promoter drives gene expression in a set of sensory neurons and interneurons [31]. Among them, PVP and PVQ, for instance, display extensive gap junction connections [30]. Indeed, characteristic UNC-9::GFP puncta were observed on the neurites and cell bodies with this P*unc-53*::UNC-9::GFP marker (S4A Fig). Interestingly, when either *ire-1* or *xbp-1* is mutated, the UNC-9::GFP signal becomes diffusely distributed to the perinuclear region of those neurons (S4A Fig). Meanwhile, the GFP signal along neurites is significantly decreased (S4A Fig). Together, the Ire1-Xbp1 pathway is specifically required for the sub-cellular localization of ectopically expressed UNC-9 proteins in neurons.

### The chronic stress triggers the IRE-1-mediated *xbp-1* splicing

The excess UNC-9::GFP triggers an ER stress response in DD/VD neurons. We wondered whether *ire-1* and *xbp-1* regulate the subcellular localization of UNC-9::GFP in DD/VDs through their roles in the ER stress response. To address this question, we firstly examined in which subcellular compartments the UNC-9::GFP is localized in *ire-1* and *xbp-1* mutants. In wild-type animals, a few UNC-9::GFP puncta co-localize with the ER marker CYTB-5.1::mCherry (Fig 2C). In *ire-1* or *xbp-1* mutant animals, however, the diffuse UNC-9::GFP signal completely overlaps with the CYTB-5.1::mCherry signal (Fig 2C). The ER localization of UNC-9::GFP signal suggests that the ectopically expressed UNC-9::GFP proteins probably are misfolded or unfolded in *ire-1* and *xbp-1* mutants. Under ER stress conditions, the transcription of genes encoding ER chaperones is upregulated to facilitate the correct folding of ER-synthesized proteins. Therefore, we further examined the expression of P*hsp-4*::mCherry in *ire-1* or *xbp-1* mutant animals, and found that the specific induction of P*hsp-4*::mCherry in DD/VD neurons was dramatically reduced (Fig 2D and S1A Fig). We noticed that the basal expression level of the *hsp-4* reporter is also reduced by *ire-1* or *xbp-1* mutation (S1A Fig). Therefore, *ire-1* and *xbp-1* are required for both induced and basal expression of *hsp-4* gene.

The induction of ER chaperone genes relies on activation of the transcription factor XBP-1s. Next, we tested whether the excess UNC-9::GFP activates the alternative splicing of *xbp-1* mRNA. We fused a cDNA encoding mCherry in-frame with the inactive form of *xbp-1* driven by the P*unc-25* promoter (Fig 2E) [32]. In the absence of ER stress, the regulatory intron cannot be spliced out, which results in “mCherry on”. In contrast, the ER stress response will trigger the alternative spicing of *xbp-1* mRNA and the regulatory intron can be removed, resulting in “mCherry off”. When UNC-9::GFP was overexpressed in DD/VDs, we found that the “mCherry off” phenotype was induced in DD/VDs (Fig 2F). In the absence of *ire-1*, the “mCherry off” phenotype was strongly suppressed and the mCherry signal reappeared in DD/VDs (Fig 2F). Together, these results provide evidence that UNC-9::GFP overexpression in DD/VDs triggers IRE-1-mediated *xbp-1* splicing.

### Age-dependent requirement for the IRE-1-XBP-1 UPR branch under stress conditions

Utilizing the distinct UNC-9::GFP localization pattern, we explored at which stage the *ire-1-xbp-1* branch is required for the stress response triggered by excess UNC-9::GFP. From the embryonic stage to larval stage 3 (L3), we found that the majority of UNC-9::GFP signal in *ire-1* mutants displays a punctate pattern similar to wild type (Fig 3A), which suggests that the *ire-1-*mediated stress response is not required for the correct folding of excessive UNC-9 protein during this period. However, when *ire-1* animals progress to the L4 stage, the UNC-9::GFP signal on the neuronal processes is significantly decreased (Fig 3A). Meanwhile, more UNC-9::GFP signal became diffusely distributed in the cell body region (Fig 3A). Thus, the absence of the stress response has no obvious effect on the subcellular localization of ectopically expressed UNC-9::GFP until the late developmental stage.

**Fig 3.**
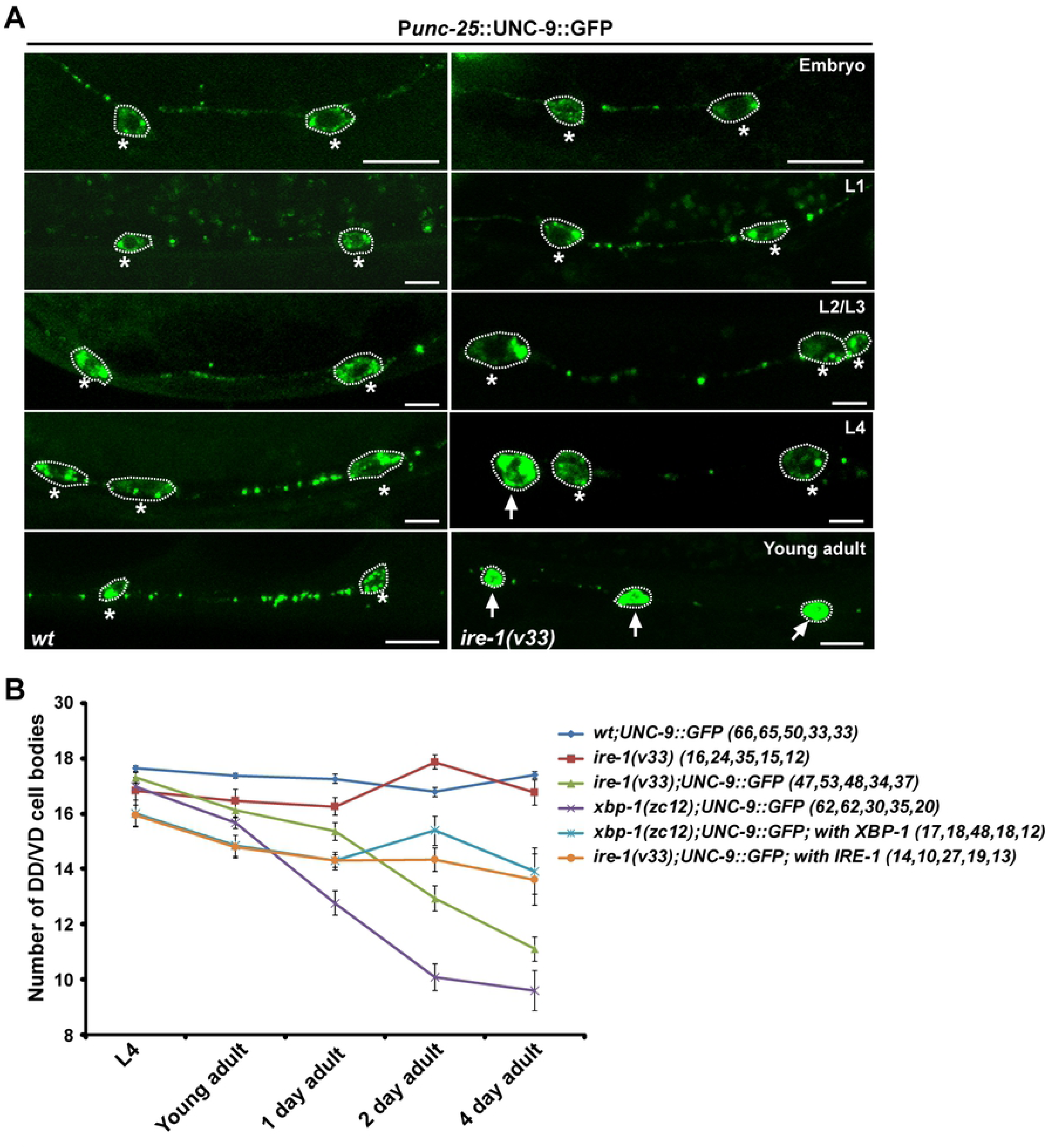
The cell-protective role of the IRE-1-XBP-1 UPR branch under stress conditions. (A) The progressive UNC-9::GFP (green) localization defect in *ire-1(v33)* animals compared with wild type. Scale bars represent 5 µm. Dashed lines encircle the DD/VD cell bodies. Individual DD/VD cell bodies are encircled by dashed lines. Asterisks indicate normal cells. White arrows point to defective cells. (B) Quantification of DD/VD cell numbers in various genotypes. *N* is indicated for each genotype.

Next, we examined whether prolonged failure of the UPR in DD/VD neurons leads to cell death. With the P*unc-25*::mCherry marker, around 17 DD/VD neurons could be unambiguously identified in wild-type animals (Fig 3B). A similar number of DD/VDs was observed when excess UNC-9::GFP was present (Fig 3B), which suggests that the functional UPR system is able to keep cells alive under ER stress conditions. In contrast, when *ire-1* or *xbp-1* was removed, the total number of DD/VDs dropped significantly (Fig 3B). Notably, the reduced cell number occurs only under stress conditions. Neither *ire-1* nor *xbp-1* mutation caused neuronal loss in the absence of excess UNC-9::GFP (Fig 3B). Introducing a wild-type copy of the *ire-1* or *xbp-1* gene into DD/VDs significantly rescued the neuronal loss in *ire-1* or *xbp-1* mutant animals respectively (Fig 3B). Hence, the IRE1-XBP1-mediated stress response has a beneficial effect during prolonged chronic stress *in vivo*.

### *pmk-3* is required for the ER stress response

In addition to *xbp-1(xd131)*, the unbiased genetic screen also identified a *pmk-3* allele, *xd74*. In *xd74* animals, the UNC-9::GFP signal is diffusely distributed in the cell body region of DD/VDs and is greatly reduced in DD/VD neuronal processes (Fig 4A), which mimics the *ire-1* or *xbp-1* mutant phenotype. Whole-genome sequencing identified a G-to-A mutation which affects the splicing donor site of the fourth intron in the *pmk-3* gene (S5A Fig). RT-PCR combined with sequencing analysis further showed that the *xd74* mutation leads to two distinct splicing alterations (S5B Fig), which both result in a premature stop codon in the *pmk-3* gene product. Other *pmk-3* alleles, including *pmk-3*(*ok169)* and *pmk-3(tm745)*, display similar UNC-9::GFP localization defects in DD/VD neurons to *pmk-3(xd74)* animals (Fig 4A and 4B). Introducing a wild-type copy of *pmk-3* into DD/VDs using the P*unc-25* promoter greatly rescued the UNC-9::GFP distribution defect in *pmk-3* mutant animals (Fig 4A and 4B), which indicates that loss of function of *pmk-3* is responsible for the UNC-9::GFP mis-localization. *pmk-3* encodes one of the three p38 MAPKs in *C. elegans* and the kinase domain is essential for PMK-3 function in UNC-9::GFP localization in DD/VDs (Fig 4A and 4B). Because animals carrying a single mutation of any of the other MAPK components do not show any obvious UNC-9::GFP distribution abnormality (S1 Table), we concluded that p38/PMK-3 alone plays an important role in the protein localization of ectopic UNC-9::GFP in DD/VD neurons.

**Fig 4.**
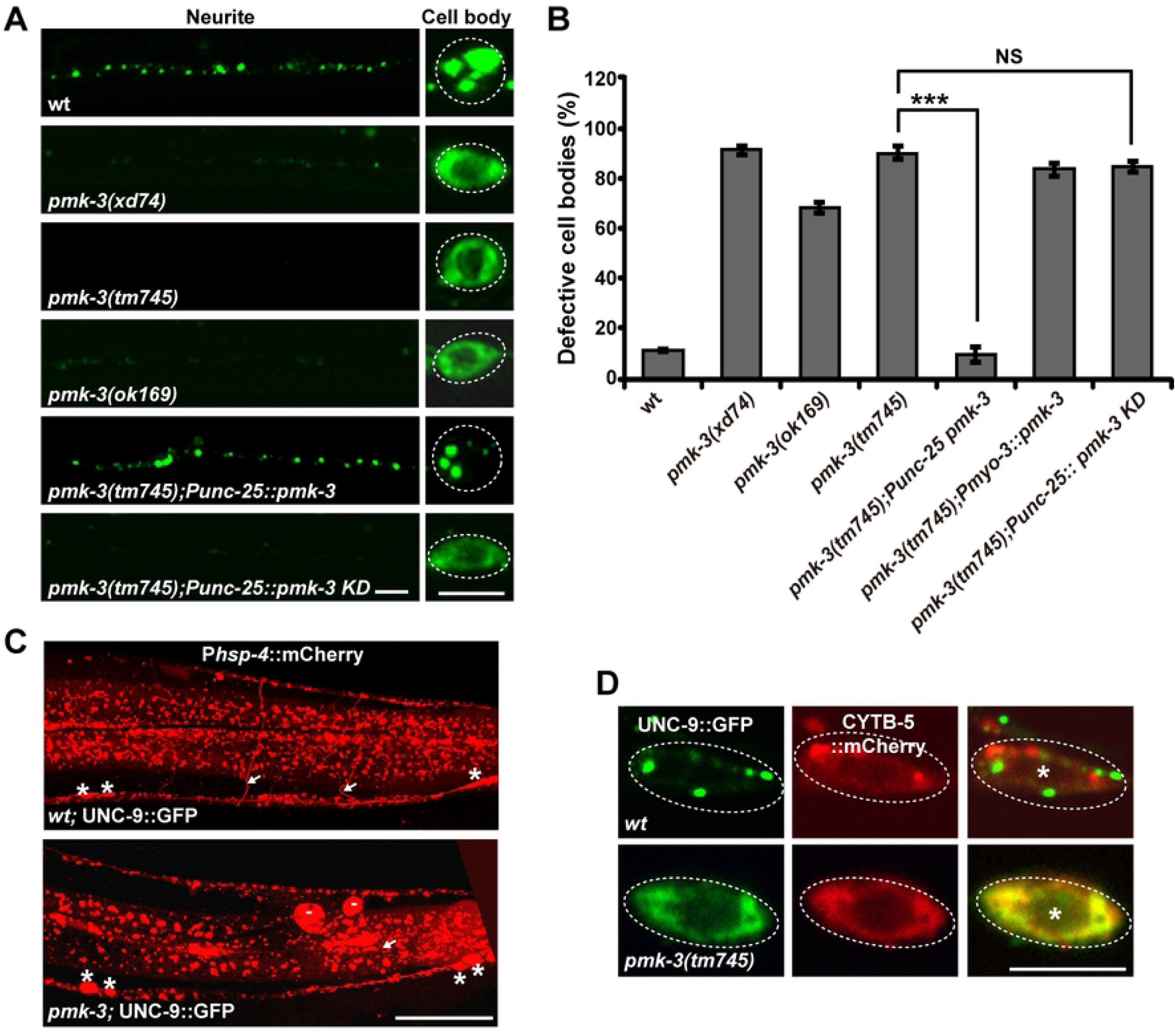
*pmk-3* is required for UPR triggered by chronic stress. (A) The localization of UNC-9::GFP (green) in various genotypes. Scale bars represent 5 µm. (B) Quantification of the UNC-9::GFP localization defect in various genotypes. *N* = 20 for each genotype; \*\*\**P* < 0.001; NS, not significant. One-way ANOVA with Dunnett’s test. (C) Compared to wild type (*wt*), expression of P*hsp-4*::mCherry (red) in the commissure region of DD/VD neurons is reduced by *pmk-3* mutation. White arrows indicate DD/VD commissures. Asterisks indicate DD/VD cell bodies. Scale bar represents 25 µm. (D) Co-localization analysis between UNC-9::GFP (green) and the ER marker CYTB-5::mCherry (red) in the DD or VD cell body region of wild-type and *pmk-3* animals. Asterisks indicate DD/VD cell bodies. Scale bar represents 5 µm.

*pmk-3* mutation does not affect the punctate distribution of UNC-9::GFP at the cell-cell junction region in muscle cells (S3A Fig). Therefore, *pmk-3* is probably not involved in the subcellular localization of UNC-9 protein in general. The similarity between *pmk-3* and *ire-1* or *xbp-1* mutant animals strongly hinted that *pmk-3* is also involved in mediating the stress response triggered by UNC-9::GFP-overexpression. To confirm this, we examined the induction of the P*hsp-4*::mCherry reporter in DD/VDs. As shown in Fig 4C, most of the 19 commissures of DD/VD neurons were easily detected with P*hsp-4*::mCherry in wild-type animals, while only 1 or 2 DD/VD commissures were seen in *pmk-3* mutants. We noticed that the individual DD/VD cell bodies were still identified with P*hsp-4*::mCherry in *pmk-3* mutants (Fig 4C), which suggests that the stress response is partially affected by *pmk-3*. We further examined the UNC-9::GFP distribution and found that the diffuse UNC-9::GFP signal in *pmk-3* mutants completely overlaps with the ER marker CYTB-5.1::mCherry in DD/VD cell body regions (Fig 4D). This suggests that misfolded or unfolded UNC-9::GFP proteins have accumulated in the ER of these cells.

### PMK-3 regulates the phosphorylation status of IRE-1

The excessive UNC-9::GFP triggers the *ire-1*-*xbp-1*-mediated stress response. How does PMK-3 participate in this process? Firstly, we addressed the genetic interactions between *pmk-3*, *ire-1* and *xbp-1*. In *pmk-3(tm745);ire-1(v33)* and *pmk-3(tm745);xbp-1(zc12)* double mutants, the UNC-9::GFP distribution resembles the *ire-1* or *xbp-1* null (Fig 5A and S6A Fig). The *pmk-3(tm745);ire-1(v33); xbp-1(zc12)* triple mutant also mimics *ire-1* or *xbp-1* single null mutants (Fig 5A and S6A Fig), which is in agreement with the notion that *pmk-3* functions in the same process with *ire-1* and *xbp-1*. Intriguingly, when the wild-type *ire-1* gene was overexpressed in *pmk-3* mutants, the UNC-9::GFP distribution defect was efficiently rescued (Fig 5A and S6A Fig). In contrast, neither *ire-1* nor *xbp-1* mutant phenotype was suppressed by *pmk-3* overexpression (Fig 5A and S6A Fig). IRE-1 acts on XBP-1 activation to regulate the stress response. We further found that overexpression of the active but not the inactive form of *xbp-1* also rescued the *pmk-3* mutant phenotype (Fig 5A and S6A Fig). The suppression of the *pmk-3* mutant phenotype by *ire-1* was totally lost when the *xbp-1* gene was removed (Fig 5A and S6A Fig). In addition, the alternative splicing of *xbp-1* mRNA in DD/VD cells is partially affected by *pmk-3* mutation (S6B Fig). Together, these results indicate that *pmk-3* functions through the *ire-1*-*xbp-1* branch to regulate the stress response.

**Fig 5.**
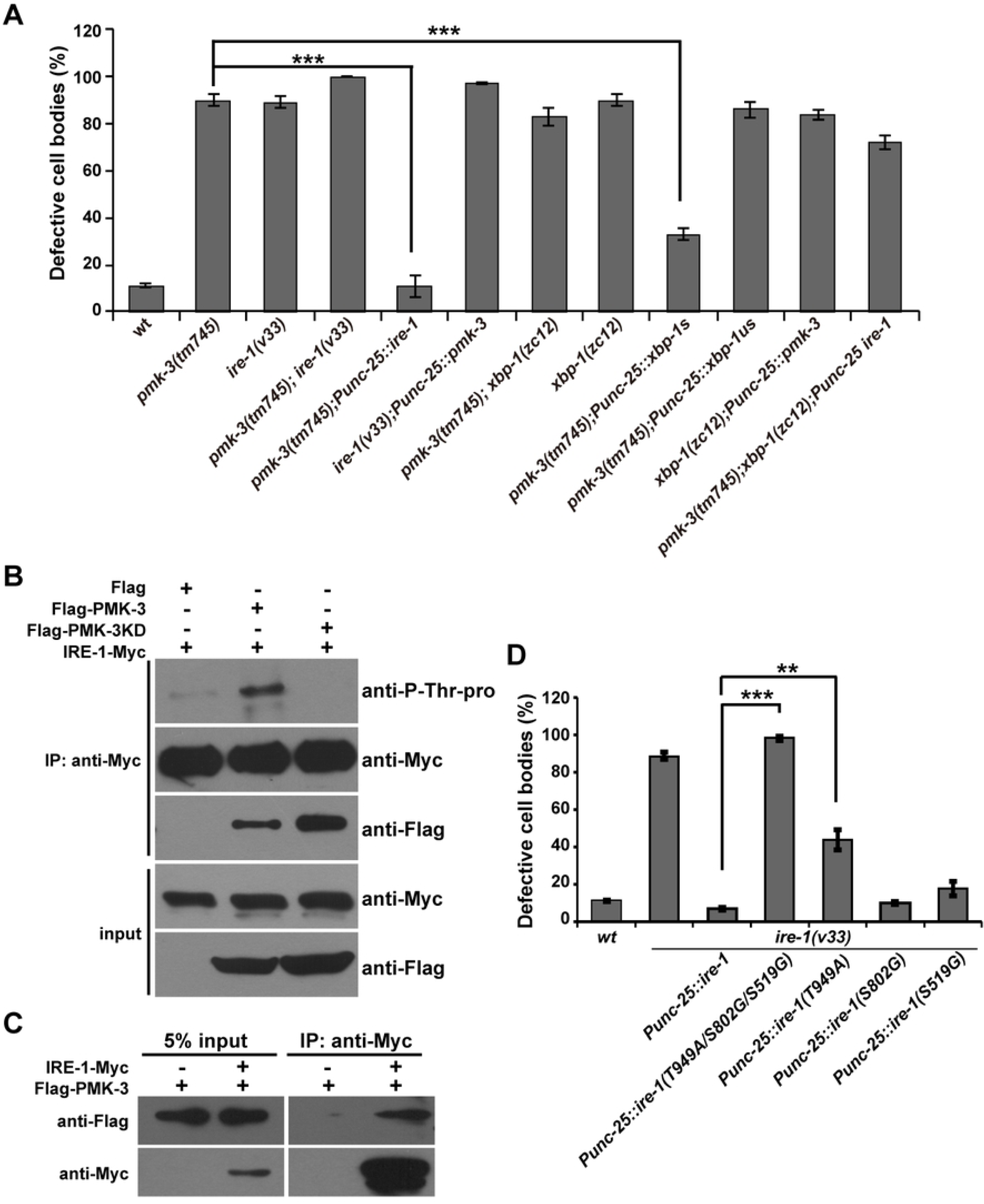
PMK-3 regulates the phosphorylation status of IRE-1. (A) Quantification of the UNC-9::GFP localization defect in various genotypes. *N* = 20 for each genotype; \*\*\**P* < 0.001; NS, not significant. One-way ANOVA with Dunnett’s test. (B) The phosphorylation of IRE-1-Myc is enhanced in the presence of Flag-PMK-3 but not Flag or Flag-PMK-3KD (KD: kinase dead). (C) IRE-1-Myc is precipitated by Flag-PMK. (D) Quantification of the rescue activity of various *pmk-3* mutations. *N* = 20 for each genotype; \*\*\**P* < 0.001; \*\**P* < 0.01; NS, not significant. One-way ANOVA with Dunnett’s test.

We showed above that the kinase-defective (KD) mutant form of PMK-3 (*pmk-3* KD) lost its capability to rescue the UNC-9::GFP localization defect in *pmk-3* mutants (Fig 4A and 4B). This suggests that the kinase activity of PMK-3 is crucial for its function. The cytosolic region of IRE-1 contains three putative p38 MAPK phosphorylation sites: Ser519 (S519), Ser802 (S802) and Thr949 (T949). T949 is located on the C-terminal RNase domain of IRE-1 and phosphorylation of the corresponding site in human (T973) contributes to the full RNase activity of Ire1 *in vivo* [33]. To test whether PMK-3 regulates the stress response through phosphorylation of IRE-1, we performed the following experiments. Firstly, we tested whether the presence of PMK-3 altered the phosphorylation level of IRE-1. By incubating IRE-1 protein with either the wild-type or kinase-inactive form of PMK-3, we found that the wild-type but not the KD PMK-3 enhanced the phosphorylation of IRE-1 (Fig 5B). Second, we tested whether PMK-3 can associate with IRE-1. After affinity purification, we found that the Flag-tagged PMK-3 protein was co-immunoprecipitated with IRE-1-Myc (Fig 5C), which is suggestive of direct binding between PMK-3 and IRE-1. Catalytically inactive MAP kinases are often used to detect interactions between MAP kinases and their substrates or interacting partners [34]. Indeed, the constitutively inactive form of PMK-3 associated with IRE-1 with even higher affinity (Fig 5B). Third, we mutated the S519, S802 or T949 residues individually to alanine or glycine and found that the T949A substitution significantly decreased the rescuing activity of IRE-1 (Fig 5D). Further mutating the S519 and S802A sites (S519G/S802G/T949A triple mutation) completely blocked the rescuing activity of PMK-3 (Fig 5D). Together, these results provide evidence that PMK-3 phosphorylates IRE-1 to regulate the ER stress response.

### *daf-2* suppresses the ER stress response defect in *pmk-3*, *ire-1* and *xbp-1* mutants

Considering that ER stress underlies many pathological conditions, we utilized the *xbp-1(xd131)* strain and performed a further genetic suppressor screen to explore whether the malfunctioning ER stress response can be repressed. From the screen, we identified the *xd211* mutation. In the presence of *xd211*, the punctate pattern of UNC-9::GFP was restored along the neurites and in the cell body region of DD/VD neurons (Fig6A and 6B). Genetic mapping and whole-genome sequencing identified a threonine-to-lysine change at codon 926 in the insulin/IGF-1 receptor DAF-2. The UNC-9::GFP distribution defect of *xbp-1(xd131)* or *xbp-1(zc12)* was also suppressed by the *daf-2(e1370)* allele (Fig 6A and 6C) (Materials and methods). This suggests that the loss of *daf-2* function is responsible for the genetic suppression of *xbp-1*. DAF-2 negatively regulates the FOXO transcription factor DAF-16. In *xbp-1(xd131) daf-2(xd211);daf-16(mu86)* triple mutant animals, the suppression of *xbp-1* by *daf-2* is completely blocked by *daf-16* mutation (Fig 6A and 6B), which indicates that the suppressive effect of *daf-2* relies on *daf-16*. Both *ire-1* and *pmk-3* function through *xbp-1*. We found that the UNC-9::GFP localization defect caused by *pmk-3* was suppressed by *daf-2* (Fig 6A and 6C). To a lesser degree, the diffuse UNC-9::GFP localization phenotype in *ire-1* was also partially suppressed by *daf-2* (Fig 6A and 6C). Thus, the insulin pathway functions antagonistically with the *pmk-3*-*ire-1*-*xbp-1* pathway to regulate the stress response (S7A Fig).

**Fig 6.**
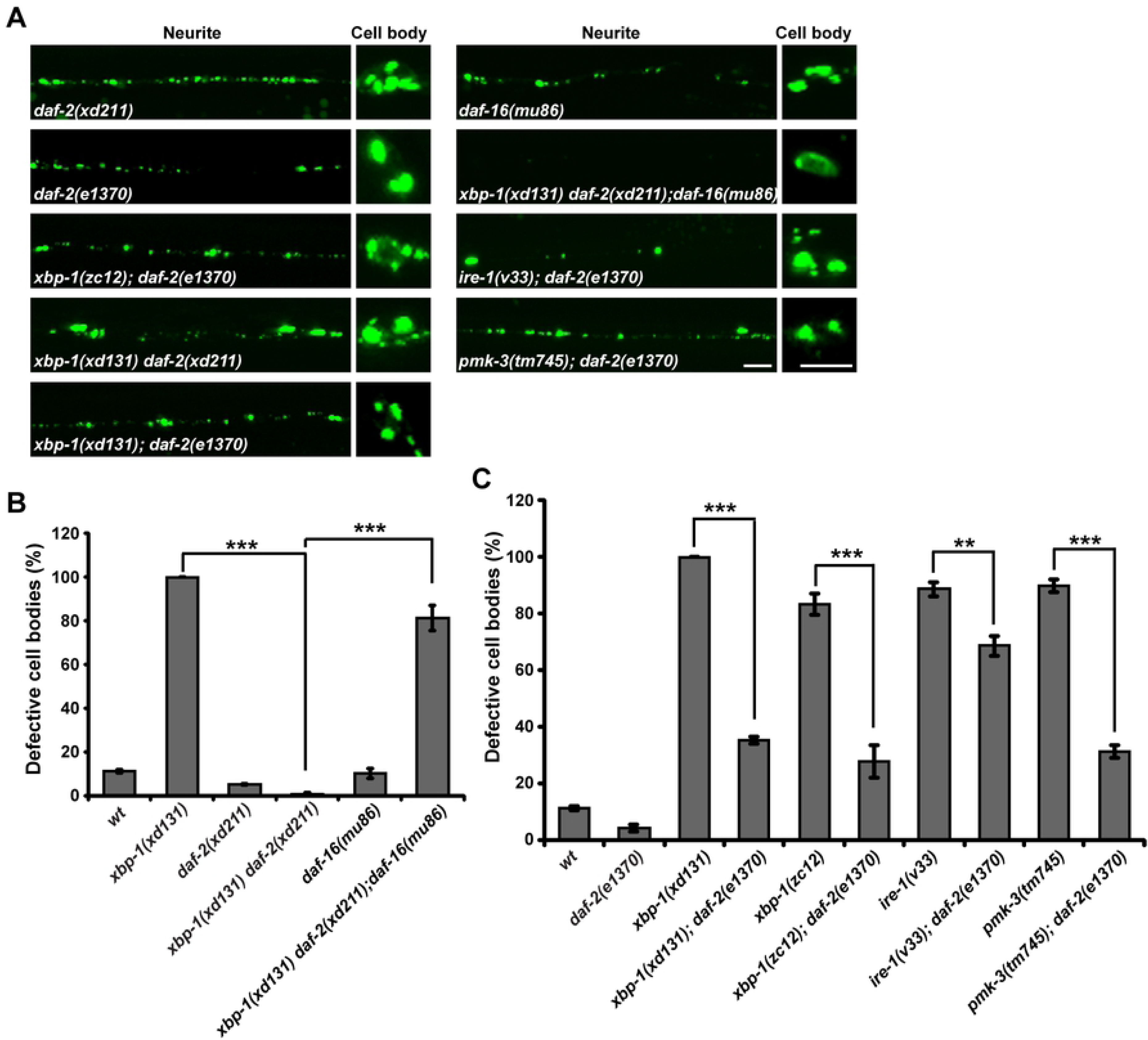
*daf-2* suppresses *pmk-3*, *ire-1* and *xbp-1* mutant phenotype. (A) The localization of UNC-9::GFP (green) in various genotypes. Scale bars represent 5 µm. (B) Quantification of the UNC-9::GFP localization defect in various genotypes. *N* = 20 for each genotype; \*\*\**P* < 0.001; One-way ANOVA with Dunnett’s test. (C) Quantification of the UNC-9::GFP localization defect in various genotypes. *N* = 20 for each genotype; \*\*\**P* < 0.001; \*\**P* < 0.01. One-way ANOVA with Dunnett’s test.

### *daf-2* restores autophagy induction in *pmk-3*, *ire-1* and *xbp-1* mutant animals

How is the suppression effect of *daf-2* on *pmk-3*, *ire-1* and *xbp-1* achieved? To answer this, we firstly examined induction of the ER chaperone reporter P*hsp-4*::mCherry. In *daf-2* single mutant animals, the intensity of P*hsp-4*::mCherry signal is indistinguishable from wild type, which suggests that the loss of *daf-2* function does not obviously alter *hsp-4* gene induction (Fig 7A). The strong reduction of *hsp-4* expression previously observed in *xbp-1* or *ire-1* mutants was not rescued in *daf-2;xbp-1* or *daf-2;ire-1* double mutants (Fig 7A). Similarly, the partial reduction of *hsp-4* expression in *pmk-3* mutants was not restored by *daf-2* (Fig 7B). As shown in Fig 7B, the reduced P*hsp-4*::mCherry signal on DD/VD neuron processes in *pmk-3* mutants was not rescued by further removal of *daf-2* (Fig 7B). In addition, overexpressing *hsp-4* in DD/VDs did not rescue the UNC-9::GFP localization defect in *ire-1* or *xbp-1* mutant animals (S7B Fig).

**Fig 7.**
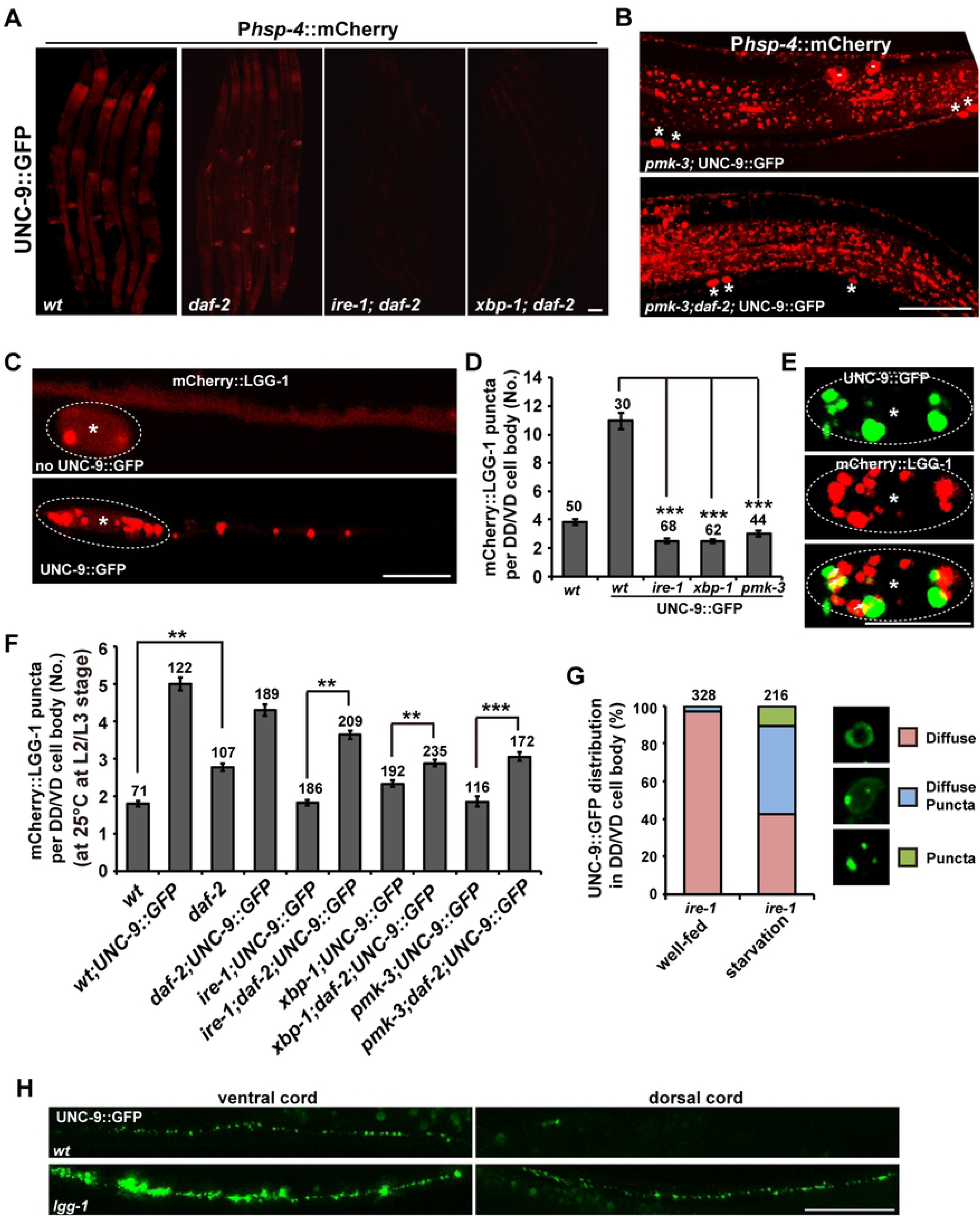
Autophagy induction is restored by *daf-2* in *pmk-3*, *ire-1* and *xbp-1* mutants. (A) The reduction of P*hsp-4*::mCherry expression (red) in *ire-1* or *xbp-1* mutants is not rescued by *daf-2*. (B) The partial reduction of P*hsp-4*::mCherry expression (red) in *pmk-3* mutants is not suppressed by *daf-2*. Asterisks indicate DD/VD cell bodies. Scale bars in (A) and (B) represent 25 µm. (C) The distribution of mCherry::LGG-1 (red) in the absence or presence of UNC-9::GFP in DD/VD neurons. Scale bar represents 5 µm. (D) Quantification of the number of mCherry::LGG-1 puncta in DD/VD cell body regions. *N* is indicated for each genotype. (E) The UNC-9::GFP (green) puncta partially co-localize with mCherry::LGG-1 (red). Scale bar represents 5 µm. Asterisks indicate DD/VD cell bodies. (F) Quantification of the number of mCherry::LGG-1 puncta in DD/VD cell body regions in various genotypes. *N* is indicated for each genotype. (G) Starvation treatment partially suppresses the *ire-1* mutant phenotype. The different UNC-9::GFP (green) distribution patterns in DD/VD cell body region are indicated by the colored boxes. *N* is indicated for each genotype. (H) The UNC-9::GFP distribution in wild-type and *lgg-1* mutant animals. Scale bar represents 25 µm.

Autophagy is a catabolic mechanism that delivers misfolded proteins and damaged organelles for degradation. We next asked whether autophagy is involved in *daf-2* suppression. We firstly examined whether overexpression of UNC-9::GFP induced autophagy in DD/VDs. Using mCherry-tagged LGG-1 (*C. elegans* ortholog of the yeast autophagy protein Atg8) driven by the *unc-25* promoter, we examined the autophagy activity in DD/VD neurons. In the absence of UNC-9::GFP, mCherry::LGG-1 is diffusely distributed along the neuronal processes and there are typically less than 4 mCherry::LGG-1 puncta in the cell body region (Fig 7C and 7D). When UNC-9::GFP was overexpressed, the number of mCherry::LGG-1 puncta dramatically increased in both the cell body region (more than 10 puncta) and the neuronal processes of DD/VD neurons (Fig 7C and 7D), which suggests that autophagy was induced. In addition, the excess UNC-9::GFP puncta are closely adjacent to or partially overlapping with the mCherry::LGG-1 signal (Fig 7E). This indicates that the excess UNC-9 protein may undergo autophagy-mediated degradation. In the absence of *pmk-3*, *ire-1* or *xbp-1*, the autophagy induction in DD/VDs was dramatically decreased (Fig 7D), which suggests that the p38-IRE1-XBP1 pathway is required for autophagy induction triggered by excess UNC-9::GFP.

We then tested whether *daf-2* could suppress the *pmk-3*, *ire-1* or *xbp-1* mutant phenotypes by acting through autophagy induction. In the absence of excess UNC-9::GFP, the basal level of autophagy in DD/VD cells is slightly increased by *daf-2* mutation (Fig 7F). In the presence of UNC-9::GFP, the autophagy induction in DD/VD cells in *daf-2* mutants remains at a similar level to that in wild type (Fig 7F). Intriguingly, when the *daf-2* mutation was introduced into either *ire-1*, *xbp-1* or *pmk-3* mutants, the number of mCherry::LGG-1 puncta in DD/VD neurons is significantly increased compared to the *ire-1*, *xbp-1* or *pmk-3* single mutants (Fig 7F). Because the reduction of autophagy activity in *ire-1* or *xbp-1* or *pmk-3* mutant animals could be counteracted by loss of *daf-2* function, we wondered whether increasing the autophagy activity alone could suppress the UPR failure caused by the defective *pmk-3*-*ire-1*-*xbp-1* pathway. Starvation treatment activates autophagy [35, 36]. In well-fed *ire-1* mutants, most of the DD/VD cell bodies (more than 95%) contain diffusely distributed UNC-9::GFP signal. After starvation, the UNC-9::GFP adopted a punctate distribution pattern in a significant proportion (about 58%) of DD/VD cells (Fig 7G). This implies that starvation-activated autophagy suppresses the UPR failure in *ire-1* mutant animals. Our attempts to introduce autophagy mutations into either *daf-2 xbp-1* or *daf-2;ire-1* animals failed, probably because disturbed autophagy has a severe effect on the viability of these double mutants. However, when the autophagy process was disrupted by *lgg-1* mutation, we found that the number and intensity of UNC-9::GFP puncta was significantly increased (Fig 7H). Taken together, these results provide evidence that autophagy activation may contribute to the suppression effect of *daf-2* on the defective *pmk-3-ire-1-xbp-1*-mediated ER stress response.

## Discussion

Accumulation of exogenous or abnormal misfolded proteins in neurons contributes to the pathology associated with neurodegeneration. Our understanding of the molecular events linking ER stress to neurodegeneration requires deeper knowledge of the mechanisms that underlie ER homeostasis, including how gene transcription is activated and autophagy is regulated in specific neuronal cells *in vivo*.

In response to ER stress, transcription of the ER chaperone gene is activated by the IRE-1/XBP-1 pathway [37]. Indeed, when worms are exposed to tunicamycin or dithiothreitol (DTT), or to global protein misfolding caused by heat shock, a GFP reporter driven by the *hsp-4* promoter is strongly induced in a wide range of tissues and cells [9, 38]. Pan-neuronal ER stress also triggers *hsp-4* expression cell-non-autonomously in gut and hypodermis [39]. In contrast to previous observations though, UNC-9::GFP overexpression in DD/VD motor neurons induces ER stress only within DD/VD cells. *C. elegans* contains 302 neurons and the P*unc-25* promoter limits UNC-9::GFP overexpression to less than 30 neurons (mostly DD and VD neurons). Although the detailed mechanisms underlying this restricted effect remain elusive, it is possible that the broadness or/and the severity of ER stress in the nervous system determines whether the stress response can spread to neighboring tissues.

The cell-autonomous stress response triggered by overloading of UNC-9::GFP protein ensures that UNC-9::GFP is correctly folded and thus adopts its distinctive punctate pattern in DD/VD neurons. In contrast, when UPR fails, such as in *ire-1* or *xbp-1* mutants, a diffuse UNC-9::GFP signal accumulates in the ER. Evidently, with no need for drug treatment or any other external stimuli, the distinct UNC-9::GFP subcellular localization offers a dual marker to monitor functional verse defective UPR at single-cell resolution in an intact animal. Notably, the subcellular localization of endogenous UNC-9 and of ectopic muscle-expressed UNC-9::GFP is unaffected by *ire-1* or *xbp-1*. Aided by unbiased genetic screens, we further identified that *pmk-3*/p38 MAPK is specifically involved in the ER stress response triggered by excess UNC-9::GFP. Among the three p38 MAPKs in *C. elegans* [40], PMK-1 is extensively involved in innate immunity and subsequently in DAF-2 or/and DAF-16-related longevity [41–46]. While the *ire-1*-*xbp-1* pathway protects animals from *Pseudomonas aeruginosa* infection, inactivation of the PMK-1-mediated immune response suppresses the lethal effect of P. *aeruginosa* infection in *xbp-1* mutants [47]. Here, in the context of UNC-9::GFP overexpression, *pmk-3* functions similarly to *ire-1* and *xbp-1*. In contrast, neither PMK-1 nor the upstream MAPKK (MAPK kinase) SEK-1 nor MAPKKK (MAPK kinase kinase) NSY-1 [46] is required for UNC-9::GFP-induced UPR (Supplementary Table 1). What determines the differential roles of p38 MAPKs in various biological processes? Compared to the wide range of reactions from multiple tissues during pathogen infection, we suspect that protein overloading in a handful of neurons triggers a rather mild stress response. Although the core components (for instance, *ire-1* and *xbp-1*) are shared, other regulatory components are employed in a tissue- or/and intensity-dependent manner. How does PMK-3/p38 participate in this cell-autonomous chronic ER stress response? A previous study showed that p38 MAPK directly phosphorylates the activated form of XBP1 and enhances its nuclear migration [48]. Here, however, we found that overexpressing *ire-1* suppresses the *pmk-3* mutant phenotype, while upregulation of *pmk-3* does not bypass the functional requirement for *ire-1*. This suggests that *pmk-3* may function upstream of *ire-1*. Consistent with this idea, PMK-3 binds to IRE-1 and the phosphorylation status of IRE-1 is affected by the presence of functional PMK-3. Among three putative p38 MAPK phosphorylation sites in IRE-1, T949 is located in the C-terminal RNase domain. Phosphorylation of the corresponding site does not have a direct impact on the splicing activity *in vitro*, but has a noticeable effect *in vivo* [33]. Here, we showed that the alternative splicing of *xbp-1* mRNA in DD/VD neurons is partially affected by *pmk-3* and mutation of this T949 site significantly decreased the rescue activity of IRE-1. Given that phosphorylation on the RNase domain may be involved in the recruitment of unknown cofactors to maximize the RNase activity of IRE-1 [33], it is fascinating to speculate that the regulatory role of PMK-3/p38 MAPK is particularly important in fighting against chronic ER stress caused by gradual accumulation of mutated or overexpressed protein, which is common under pathogenic conditions in many human diseases.

The relationship between insulin signaling and UPR regulation is rather complicated [49–51]. Worms with loss-of-function mutations in the insulin/IGF-1 receptor *daf-2* are long-lived, and both *ire-1* and *xbp-1* make a large contribution to the long lifespans of *daf-2* mutants and also increase their ER stress resistance [35, 52]. The *daf-2* mutation also improved the ER stress resistance of *xbp-1* and *ire-1* mutant animals when they were treated with low but not high tunicamycin concentrations, and DAF-16 acts through ER-associated degradation systems independently of *ire-1* [53]. A more recent study further showed that the complexity of PVD neuron dendrite arborization is greatly reduced by a loss-of-function mutation in *ire-1*, and further reducing insulin signaling restored the normal PVD neuronal morphology [54]. However, *xbp-1* is not involved in PVD morphogenesis at all, which suggests that the antagonistic role of insulin signaling may not act through the non-canonical splicing of *xbp-1* mRNA as in the UPR process [54]. Here, we found that *daf-2* mutation suppresses the defective ER stress response in both *ire-1* and *xbp-1* animals and this suppression is dependent on the FOXO transcription factor DAF-16. Here, the cell autonomous reduction of *hsp-4* expression in *ire-1* or *xbp-1* mutants is not restored by *daf-2*. Thus, DAF-16 may act on alternative path to regulate the stress response.

Intriguingly, autophagy induction was restored in *ire-1* or *xbp-1* mutant cells by loss of *daf-2* function. It has been shown that *daf-2* mutants exhibit increased levels of autophagy, and autophagy is required for their extended lifespan [36]. In addition, the basal level of autophagy is increased by *daf-2* mutation in both wild-type and *ire-1* mutant animals [53,55,56]. Here, we found that although the excess UNC-9-triggered autophagy up-regulation in DD/VD neurons is not further enhanced by *daf-2* mutation, the reduction of autophagy in *ire-1* or *xbp-1* mutants was indeed suppressed by *daf-2*. The FOXO forkhead transcription factors are key transcriptional regulators of autophagy [57]. Indeed, the autophagy induction by excess UNC-9::GFP is completely suppressed by *daf-16* mutation (data not shown). We suspected that in *ire-1* or *xbp-1* single mutants, while XBP-1 activation is defective, the DAF-16/FOXO transcription factor remains inactive due to the presence of DAF-2. In these single mutants, neither transcription of ER chaperone genes (for instance *hsp-4*/Bip) nor autophagy induction occurs, and the proper folding of excess protein could not be achieved. When *daf-2* is removed, the DAF-16 transcription factor can be activated independent of *xbp-1*. Thus, although the transcriptional upregulation of ER chaperone genes (such as *hsp-4*/Bip) may be missing, active DAF-16 turns on autophagy, which may act through protein degradation or ER homeostasis to facilitate protein folding in *ire-1* and *xbp-1* mutants. The accumulation of misfolded proteins in the ER appears not to directly activate DAF-16 [53]. Therefore, DAF-16-mediated autophagy is crucial only under conditions in which both DAF-2/insulin receptor and the *ire-1*-*xbp-1* branch of the UPR are removed. Nevertheless, the availability of two parallel systems (S7A Fig) suggests that induction of autophagy using small molecules may be an alternative approach to treat ER stress response failure in many degenerative diseases.

## Materials and Methods

### *C. elegans* genetics

Culture and manipulation of *C. elegans* strains were performed using standard methods [58]. Mutants used in this study are listed here: LGI: *dlk-1(ju476),dlk-1(tm4024),daf-16(mu86),mom-4(ne1539),mtk-1(ok1382).* LGII: *ire-1(v33), nsy-1(ok593), xdIs15*(P*unc-25*::UNC-9::GFP;P*odr-1*::GFP). LGIII: *xbp-1(zc12),xbp-1(xd131),xbp-1(tm2457),xbp-1(tm2482), daf-2(e1370 ts), daf-2(xd211), kin-18(pk71), xbp-1(zc12),.* LGIV: *pmk-3(tm745)*, *pmk-3(ok169)*, *pmk-3(xd74)*, *jnk-1(gk7)*, *kgb-1(um3)*, *pmk-1(km25)*, *kgb-2(gk361)*, *unc-30(ju54)*, *mak-2(tm2927)*, *wnk-1(ok266), lgg-1(bp500), xdIs13*(P*unc-25*::UNC-9::GFP;P*odr-1*::GFP). LGV: *rpm-1(ju44)*, *mlk-1(ok247).* LGX: *atf-6(ok551)*, *atf-6(tm1153)*, *pek-1(ok275)*, *pek-1(ok275)*, *mkk-4 (ok1545)*, *jkk-1(km2)*, *mek-1(ks54)*, *sek-3(tm1344)*, *sek-6(tm4305 tm4136)*, *sek-1(km4).* The genetic screen was performed according to the previous report [59]. Briefly, *xdIs15* or *xbp-1(xd131);xdIs15* worms were treated with EMS (ethylmethane sulfonate) solution for 4 hours. F2 worms were examined for phenotype alteration at the young adult stage under a fluorescence microscope. Mutant animals were recovered to fresh plates for maintenance. A total of 10,000 and 5,000 mutagenized haploid genomes were screened respectively for *xdIs15* and *xbp-1(xd131);xdIs15* worms. The *pmk-3(xd74)* and *xbp-1(xd131)* mutants were isolated from the genetic screen based on the *xdIs15* strain. The *daf-2(xd211)* mutant was isolated from the genetic screen based on suppression of the *xbp-1(xd131);xdIs15* phenotype. All isolated mutants were outcrossed with wild type at least four times. For UNC-9::GFP phenotype characterization, strains containing *daf-2(e1370 ts)* were cultured at 16°C until L2/L3 stage, then shifted to 25°C to reach adulthood before phenotypic examination. Other strains were cultured at 22°C and the phenotype was characterized at the young adult stage.

### DNA constructs and transgenic animals

The *unc-25* promoter was inserted between the *BamH*I and *Sph*I sites of the pSM-GFP vector (a generous gift from Dr. Kang Shen) and the UNC-9 cDNA was inserted into the *BamH*I site. *xbp-1*, *ire-1* and *pmk-3* cDNAs were amplified by reverse transcription. The ER stress reporter plasmid P*unc-25*::XBP-1us::mCherry was modified from XB3 plasmid (a generous gift from Dr. Christopher Rongo). P*unc-25*::*cytb-5.1*::mCherry was kindly provided by Dr. Yingchuan Qi. P*unc-25*:: mCherry::MANS, P*unc-25*:: mCherry::RAB-7, P*unc-25*:: mCherry::RAB-5, and P*unc-25*::mCherry::LGG-1 were constructed by the recombination method. Transgenic animals were produced as previously described [59]. Integrated strains were obtained by UV irradiation. The strain BCN1071 which expresses P*hsp-4*::mCherry was kindly provided by Ben Lehner. All integrated transgenic animals were out-crossed at least 3 times. The corresponding transgenes are listed in S2 Table.

### Image collection and quantification

Animals were mounted on 2% agar pads in M9 buffer containing 1% 1-phenoxy-2-propanol and examined by fluorescence microscopy unless indicated otherwise. Fluorescence photographs were taken using a Zeiss Axioimager A1 with an AxioCam digital camera and Axiovision rel. 4.6 software (Carl Zeiss) or an IX81 Olympus inverted confocal microscope. All images were taken at the young adult stage unless specifically indicated. The UNC-9::GFP distribution defect in the cell body region was quantified by counting the percentage of cell bodies with diffusely localized UNC-9::GFP. To quantify the number of DD/VD neurons, the P*unc-25*::mCherry marker was used. P*hsp-4*::mCherry was used to detect induction of the chaperone gene *hsp-4* in the ER stress response. To quantify the *xbp-1* splicing activity, the P*unc-25*::XBP-1us::mCherry plasmid was used and DD/VD cell bodies with mCherry signal were counted. To characterize the distribution of UNC-9::GFP under starvation conditions, well-fed L4 *ire-1(v33);xdIs15* worms were collected and washed 8 times with M9 buffer. After sedimentation, worms were plated on NGM plates without food. After 24 hours, the starved adult worms were examined using Leica TCS SP8 confocal microscopy. All graphical data are presented as mean ± s.e.m. Two-tailed unpaired and paired Student’s *t*-tests were performed for comparison between two groups of samples. To compare multiple groups, one-way ANOVA was used with an appropriate multiple comparisons *post hoc* test (the test used is stated in each figure legend). \**P* < 0.05; \*\**P* < 0.01; \*\*\**P* < 0.001; NS, not significant.

### Quantification of autophagy

To analysis autophagy activity in DD/VD neurons, P*unc-25*::mCherry::LGG-1 was constructed. Images at the young adult stage were collected and the numbers of mCherry::LGG-1 foci in DD/VD cell body regions were counted. To characterize mCherry::LGG-1 puncta in strains carrying *daf-2((e1370 ts)*, embryos were collected and raised at 25°C until L2/L3 stage before phenotypic examination.

### Immunoprecipitation and western blotting

HEK293T (human embryonic kidney) cells were maintained in Dulbecco’s modified Eagle’s medium containing 12% FBS. For cell transfection, polyethylenimine (Polysciences) was used according to the manufacturer’s instructions. The clones used for transfection were constructed in pcDNA3.1/myc-His (-) and pFLAG-CMV-2 or their modified forms. C-terminal tags, such as GST, GFP, Flag-His and so on, were cloned into pcDNA3.1/myc-His (-) by replacing the myc-His tag. 24 hours after transfection, cells were harvested and lysed for 30 min at 4°C. After centrifugation, the supernatants were incubated with anti-FLAG M2 affinity gel beads at 4°C overnight and then washed three times with washing buffer and incubated with SDS-sample buffer. Samples were resolved by standard immunoblotting techniques. All the co-immunoprecipitation experiments were repeated at least three times.

### IRE-1 phosphorylation

The Myc-tagged cytoplasmic region of IRE-1 and the Flag-tagged full-length wild-type or kinase inactive form of PMK-3 were co-expressed in H293T cells and then the IRE-1 protein was immunoprecipitated with Myc antibodies. The phosphorylation level of IRE-1 was examined with P-Thr-Pro-101antibody (Cell Signal Technology #9391), which detects the conserved phosphorylation site (S/TP) for p38 MAPK. The signal intensity of individual protein bands was analyzed with ImageJ. The relative protein level was determined by subtracting the respective FLAG control. All the co-immunoprecipitation experiments were repeated at least three times.

## Acknowledgments

We thank Drs. Zhaowen Wang, Yishi Jin, Yingchuan Qi, Christopher Rongo, Ben Lehner and Yuji Kohara, the million mutation project, the TransgeneOme Project, and the Caenorhabditis Genetics Center for providing reagents, strains, and technical support. This work was supported by the National Basic Research Program of China (2018YFA081104 and 2016YFA0501000), the National Natural Science Foundation of China (31921002) and the Chinese Academy of Sciences (XDBS010010100). The funders had no role in study design, data collection and analysis, decision to publish, or preparation of the manuscript

## Legends for Supplementary Figures

**S1 Fig. *ire-1* and *xbp-1* are required for the induced and basal expression of *hsp-4*.**

(A) Expression of P*hsp-4*::mCherry (red) in wild-type (*wt*), *ire-1* and *xbp-1* animals with or without UNC-9::GFP. Scale bar represents 25 µm. (B) P*hsp-4*::mCherry (red) is induced in UNC-9::GFP (green) expressing DD/VD neurons (white arrows). Scale bar represents 50 µm

**S2 Fig. *ire-1* and *xbp-1* are required for the sub-cellular localization of ectopic UNC-9::GFP in DD/VD neurons.**

(A) Quantification of the UNC-9::GFP localization defect in various genotypes. *N* = 20 for each genotype; \*\*\**P* < 0.001; \*\**P* < 0.01. One-way ANOVA with Dunnett’s test. (A) Quantification of the mRNA expression of P*unc-25*::UNC-9::GFP in various genotypes.

**S3 Fig. The localization of UNC-9::GFP expressed in muscle cells or in knock-in line is not altered by *ire-1* or *xbp-1*.**

(A) The localization of UNC-9::GFP (green) in muscle cells in wild-type (*wt*), *ire-1(v33)*, *xbp-1(zc12)*, and *pmk-3(tm745)* animals carrying the P*myo-3*::UNC-9::GFP transgene. White lines highlight the muscle cells. Scale bar represents 25 µm. (B) UNC-9::GFP expressed at the endogenous level from a knock-in allele (UNC-9::GFP KI) is distributed on the neuronal processes and cell surface of DD/VD neurons (asterisks) in *ire-1* (A) and *xbp-1* (B)mutants. Scale bar represents 5 µm.

**S4 Fig. *ire-1* and *xbp-1* affect the localization of ectopic UNC-9::GFP in other neurons.**

(A) The localization of UNC-9::GFP in P*unc-53* expressing neurons (green) in wild-type (*wt*), *ire-1(v33)*, and *xbp-1(zc12)* animals carrying the P*unc-53*::UNC-9::GFP transgene. White arrows indicate UNC-9::GFP puncta on neurites. Dashed lines encircle the cell bodies. Scale bars represent 5 µm.

**S5 Fig. The molecular lesions of various *pmk-3* mutants.**

(A) The structure of the *pmk-3* gene. The molecular lesions are indicated for the *ok169*, *tm745* and *xd74* alleles. Green boxes indicate exons. (B) The RT-PCR results for *pmk-3* in wild type and *pmk-3(xd74)*.

**S6 Fig. *pmk-3* functions through *ire-1*-*xbp-1* pathway.**

(A) The localization of UNC-10::GFP (green) in DD/VDs in various genotypes. Dashed lines encircle the cell bodies. Scale bars represent 5 µm. (B) Alternative splicing of *xbp-1* represented by mCherry signal in wild-type (*wt*) and *pmk-3* animals. *N* is indicated for each genotype.

**S7 Fig. The working model.**

(A) The PMK-3-IRE-1-XBP-mediated UPR acts in parallel with insulin-inhibited autophagy to alleviate chronic stress induced by excess UNC-9::GFP proteins. (B) The overexpression of *hsp-4* does not suppress the mutant phenotype of *ire-1(v33)* or *xbp-1(zc12)* animals.

## References

1. Walter P, Ron D (2011) The unfolded protein response: from stress pathway to homeostatic regulation. Science 334: 1081–1086.

2. Tabas I, Ron D (2011) Integrating the mechanisms of apoptosis induced by endoplasmic reticulum stress. Nat Cell Biol 13: 184–190.

3. Papandreou I, Denko NC, Olson M, Van Melckebeke H, Lust S, et al. (2011) Identification of an Ire1alpha endonuclease specific inhibitor with cytotoxic activity against human multiple myeloma. Blood 117: 1311–1314.

4. Carrasco DR, Sukhdeo K, Protopopova M, Sinha R, Enos M, et al. (2007) The differentiation and stress response factor XBP-1 drives multiple myeloma pathogenesis. Cancer Cell 11: 349–360.

5. Mori K (2009) Signalling pathways in the unfolded protein response: development from yeast to mammals. J Biochem 146: 743–750.

6. Cox JS, Walter P (1996) A novel mechanism for regulating activity of a transcription factor that controls the unfolded protein response. Cell 87: 391–404.

7. Mori K, Kawahara T, Yoshida H, Yanagi H, Yura T (1996) Signalling from endoplasmic reticulum to nucleus: transcription factor with a basic-leucine zipper motif is required for the unfolded protein-response pathway. Genes Cells 1: 803–817.

8. Yoshida H, Matsui T, Yamamoto A, Okada T, Mori K (2001) XBP1 mRNA is induced by ATF6 and spliced by IRE1 in response to ER stress to produce a highly active transcription factor. Cell 107: 881–891.

9. Calfon M, Zeng H, Urano F, Till JH, Hubbard SR, et al. (2002) IRE1 couples endoplasmic reticulum load to secretory capacity by processing the XBP-1 mRNA. Nature 415: 92–96.

10. Ron D, Walter P (2007) Signal integration in the endoplasmic reticulum unfolded protein response. Nat Rev Mol Cell Biol 8: 519–529.

11. Roussel BD, Kruppa AJ, Miranda E, Crowther DC, Lomas DA, et al. (2013) Endoplasmic reticulum dysfunction in neurological disease. Lancet Neurol 12: 105–118.

12. Gow A, Sharma R (2003) The unfolded protein response in protein aggregating diseases. Neuromolecular Med 4: 73–94.

13. Safra M, Henis-Korenblit S (2014) A new tool in C. elegans reveals changes in secretory protein metabolism in ire-1-deficient animals. Worm 3: e27733.

14. Mizushima N (2007) Autophagy: process and function. Genes Dev 21: 2861–2873.

15. Suzuki K, Ohsumi Y (2007) Molecular machinery of autophagosome formation in yeast, Saccharomyces cerevisiae. FEBS Lett 581: 2156–2161.

16. Chan SN, Tang BL (2013) Location and membrane sources for autophagosome formation - from ER-mitochondria contact sites to Golgi-endosome-derived carriers. Mol Membr Biol 30: 394–402.

17. Hamasaki M, Shibutani ST, Yoshimori T (2013) Up-to-date membrane biogenesis in the autophagosome formation. Curr Opin Cell Biol 25: 455–460.

18. Bernales S, McDonald KL, Walter P (2006) Autophagy counterbalances endoplasmic reticulum expansion during the unfolded protein response. PLoS Biol 4: e423.

19. Yorimitsu T, Nair U, Yang Z, Klionsky DJ (2006) Endoplasmic reticulum stress triggers autophagy. J Biol Chem 281: 30299–30304.

20. Laird DW (2010) The gap junction proteome and its relationship to disease. Trends Cell Biol 20: 92–101.

21. Xia K, Ma H, Xiong H, Pan Q, Huang L, et al. (2010) Trafficking abnormality and ER stress underlie functional deficiency of hearing impairment-associated connexin-31 mutants. Protein Cell 1: 935–943.

22. Berthoud VM, Minogue PJ, Lambert PA, Snabb JI, Beyer EC (2016) The Cataract-linked Mutant Connexin50D47A Causes Endoplasmic Reticulum Stress in Mouse Lenses. J Biol Chem 291: 17569–17578.

23. Tattersall D, Scott CA, Gray C, Zicha D, Kelsell DP (2009) EKV mutant connexin 31 associated cell death is mediated by ER stress. Hum Mol Genet 18: 4734–4745.

24. Alapure BV, Stull JK, Firtina Z, Duncan MK (2012) The unfolded protein response is activated in connexin 50 mutant mouse lenses. Exp Eye Res 102: 28–37.

25. White JG, Southgate E, Thomson JN, Brenner S (1986) The structure of the nervous system of the nematode Caenorhabditis elegans. Philos Trans R Soc Lond B Biol Sci 314: 1–340.

26. Jin Y, Jorgensen E, Hartwieg E, Horvitz HR (1999) The Caenorhabditis elegans gene unc-25 encodes glutamic acid decarboxylase and is required for synaptic transmission but not synaptic development. J Neurosci 19: 539–548.

27. Miller DM, Stockdale FE, Karn J (1986) Immunological identification of the genes encoding the four myosin heavy chain isoforms of Caenorhabditis elegans. Proc Natl Acad Sci U S A 83: 2305–2309.

28. Waterston RH (1989) The minor myosin heavy chain, mhcA, of Caenorhabditis elegans is necessary for the initiation of thick filament assembly. EMBO J 8: 3429–3436.

29. Liu Q, Chen B, Gaier E, Joshi J, Wang ZW (2006) Low conductance gap junctions mediate specific electrical coupling in body-wall muscle cells of Caenorhabditis elegans. J Biol Chem 281: 7881–7889.

30. White JG, Southgate E, Thomson JN, Brenner S (1976) The structure of the ventral nerve cord of Caenorhabditis elegans. Philos Trans R Soc Lond B Biol Sci 275: 327–348.

31. Stringham E, Pujol N, Vandekerckhove J, Bogaert T (2002) unc-53 controls longitudinal migration in C. elegans. Development 129: 3367–3379.

32. Shim J, Umemura T, Nothstein E, Rongo C (2004) The unfolded protein response regulates glutamate receptor export from the endoplasmic reticulum. Mol Biol Cell 15: 4818–4828.

33. Prischi F, Nowak PR, Carrara M, Ali MM (2014) Phosphoregulation of Ire1 RNase splicing activity. Nat Commun 5: 3554.

34. Han J, Jiang Y, Li Z, Kravchenko VV, Ulevitch RJ (1997) Activation of the transcription factor MEF2C by the MAP kinase p38 in inflammation. Nature 386: 296–299.

35. Hansen M, Chandra A, Mitic LL, Onken B, Driscoll M, et al. (2008) A role for autophagy in the extension of lifespan by dietary restriction in C. elegans. PLoS Genet 4: e24.

36. Melendez A, Talloczy Z, Seaman M, Eskelinen EL, Hall DH, et al. (2003) Autophagy genes are essential for dauer development and life-span extension in C. elegans. Science 301: 1387–1391.

37. Shen X, Ellis RE, Lee K, Liu CY, Yang K, et al. (2001) Complementary signaling pathways regulate the unfolded protein response and are required for C. elegans development. Cell 107: 893–903.

38. Urano F, Calfon M, Yoneda T, Yun C, Kiraly M, et al. (2002) A survival pathway for Caenorhabditis elegans with a blocked unfolded protein response. J Cell Biol 158: 639–646.

39. Taylor RC, Dillin A (2013) XBP-1 is a cell-nonautonomous regulator of stress resistance and longevity. Cell 153: 1435–1447.

40. Berman K, McKay J, Avery L, Cobb M (2001) Isolation and characterization of pmk-(1-3): three p38 homologs in Caenorhabditis elegans. Mol Cell Biol Res Commun 4: 337–344.

41. Troemel ER, Chu SW, Reinke V, Lee SS, Ausubel FM, et al. (2006) p38 MAPK regulates expression of immune response genes and contributes to longevity in C. elegans. PLoS Genet 2: e183.

42. Zarse K, Schmeisser S, Groth M, Priebe S, Beuster G, et al. (2012) Impaired insulin/IGF1 signaling extends life span by promoting mitochondrial L-proline catabolism to induce a transient ROS signal. Cell Metab 15: 451–465.

43. Kwon G, Lee J, Lim YH (2016) Dairy Propionibacterium extends the mean lifespan of Caenorhabditis elegans via activation of the innate immune system. Sci Rep 6: 31713.

44. Singh V, Aballay A (2006) Heat-shock transcription factor (HSF)-1 pathway required for Caenorhabditis elegans immunity. Proc Natl Acad Sci U S A 103: 13092–13097.

45. Begun J, Gaiani JM, Rohde H, Mack D, Calderwood SB, et al. (2007) Staphylococcal biofilm exopolysaccharide protects against Caenorhabditis elegans immune defenses. PLoS Pathog 3: e57.

46. Kim DH, Feinbaum R, Alloing G, Emerson FE, Garsin DA, et al. (2002) A conserved p38 MAP kinase pathway in Caenorhabditis elegans innate immunity. Science 297: 623–626.

47. Richardson CE, Kooistra T, Kim DH (2010) An essential role for XBP-1 in host protection against immune activation in C. elegans. Nature 463: 1092–1095.

48. Lee J, Sun C, Zhou Y, Lee J, Gokalp D, et al. (2011) p38 MAPK-mediated regulation of Xbp1s is crucial for glucose homeostasis. Nat Med 17: 1251–1260.

49. Piperi C, Adamopoulos C, Papavassiliou AG (2016) XBP1: A Pivotal Transcriptional Regulator of Glucose and Lipid Metabolism. Trends Endocrinol Metab 27: 119–122.

50. Sha H, He Y, Yang L, Qi L (2011) Stressed out about obesity: IRE1alpha-XBP1 in metabolic disorders. Trends Endocrinol Metab 22: 374–381.

51. Herbert TP, Laybutt DR (2016) A Reevaluation of the Role of the Unfolded Protein Response in Islet Dysfunction: Maladaptation or a Failure to Adapt? Diabetes 65: 1472–1480.

52. Henis-Korenblit S, Zhang P, Hansen M, McCormick M, Lee SJ, et al. (2010) Insulin/IGF-1 signaling mutants reprogram ER stress response regulators to promote longevity. Proc Natl Acad Sci U S A 107: 9730–9735.

53. Safra M, Fickentscher R, Levi-Ferber M, Danino YM, Haviv-Chesner A, et al. (2014) The FOXO transcription factor DAF-16 bypasses ire-1 requirement to promote endoplasmic reticulum homeostasis. Cell Metab 20: 870–881.

54. Salzberg Y, Coleman AJ, Celestrin K, Cohen-Berkman M, Biederer T, et al. (2017) Reduced Insulin/Insulin-Like Growth Factor Receptor Signaling Mitigates Defective Dendrite Morphogenesis in Mutants of the ER Stress Sensor IRE-1. PLoS Genet 13: e1006579.

55. Chang JT, Kumsta C, Hellman AB, Adams LM, Hansen M (2017) Spatiotemporal regulation of autophagy during Caenorhabditis elegans aging. Elife 6.

56. Palmisano NJ, Rosario N, Wysocki M, Hong M, Grant B, et al. (2017) The recycling endosome protein RAB-10 promotes autophagic flux and localization of the transmembrane protein ATG-9. Autophagy 13: 1742–1753.

57. Lapierre LR, Kumsta C, Sandri M, Ballabio A, Hansen M (2015) Transcriptional and epigenetic regulation of autophagy in aging. Autophagy 11: 867–880.

58. Brenner S (1974) The genetics of Caenorhabditis elegans. Genetics 77: 71–94.

59. Song S, Zhang B, Sun H, Li X, Xiang Y, et al. (2010) A Wnt-Frz/Ror-Dsh pathway regulates neurite outgrowth in Caenorhabditis elegans. PLoS Genet 6.

